# Valine restriction extends survival in a *Drosophila* model of short-chain enoyl-CoA hydratase 1 (ECHS1) deficiency

**DOI:** 10.1101/2024.08.15.608013

**Authors:** Sarah Mele, Felipe Martelli, Christopher K. Barlow, Grace Jefferies, Sebastian Dworkin, John Christodoulou, Ralf B. Schittenhelm, Matthew D.W. Piper, Travis K. Johnson

## Abstract

Short-chain enoyl-CoA hydratase 1 deficiency (ECHS1D) is a rare genetic disorder caused by biallelic pathogenic variants in the *ECHS1* gene. ECHS1D is characterised by severe neurological and physical impairment that often leads to childhood mortality. Therapies such as protein and single nutrient-restricted diets show poor efficacy, whereas development of new treatments is hindered by the low prevalence of the disorder and a lack of model systems for treatment testing. Here we report on the establishment of a *Drosophila* model of ECHS1D. Flies carrying mutations in *Echs1* (CG6543) were characterised for their physical and metabolic phenotypes, and dietary intervention to improve fly model health was explored. The *Echs1* null larvae recapitulated human ECHS1D phenotypes including elevated biomarkers (S-(2-carboxypropyl)cysteamine and 2,3-dihydroxy-2-methylbutyric acid), poor motor behaviour and early mortality, and could be rescued by expression of a human *ECHS1* transgene. We observed that both restriction of valine in isolation, or all branched-chain amino acids (BCAAs - leucine, isoleucine, and valine) together, extended larval survival, supporting the idea that reducing BCAA pathway catabolic flux is beneficial in this disorder. Further, metabolic profiling revealed substantial changes to carbohydrate metabolism, suggesting that *Echs1* loss causes widespread metabolic dysregulation beyond valine metabolism. The similarities between *Drosophila* and human ECHS1D suggest that the fly model is a valuable animal system in which to explore mechanisms of pathogenesis and novel treatment options for this disorder.

## Introduction

Short-chain enoyl-CoA hydratase 1 deficiency (ECHS1D) is a rare autosomal recessive disorder characterised by progressive neurodegeneration. Patients are typically diagnosed with Leigh syndrome, a lethal mitochondrial disease, to date, approximately 70 patients have been described.^1,2^ ECHS1D is caused by biallelic pathogenic variants in the *ECHS1* gene, which encodes a mitochondrial enzyme necessary during the catabolism of the amino acid valine and short-chain fatty acids^3^ (Figure 1). ECHS1D presents with a heterogenous clinical spectrum, ranging from a rapidly fatal neonatal course to intermittent exercise-induced dystonia.^4,5^ A deficiency in ECHS1 activity leads to abnormal accumulation of toxic valine-derived metabolites, suggesting that ECHS1D is predominantly a disorder of valine catabolism, and therefore could be treatable via diet modification.^6^

**Figure 1.**
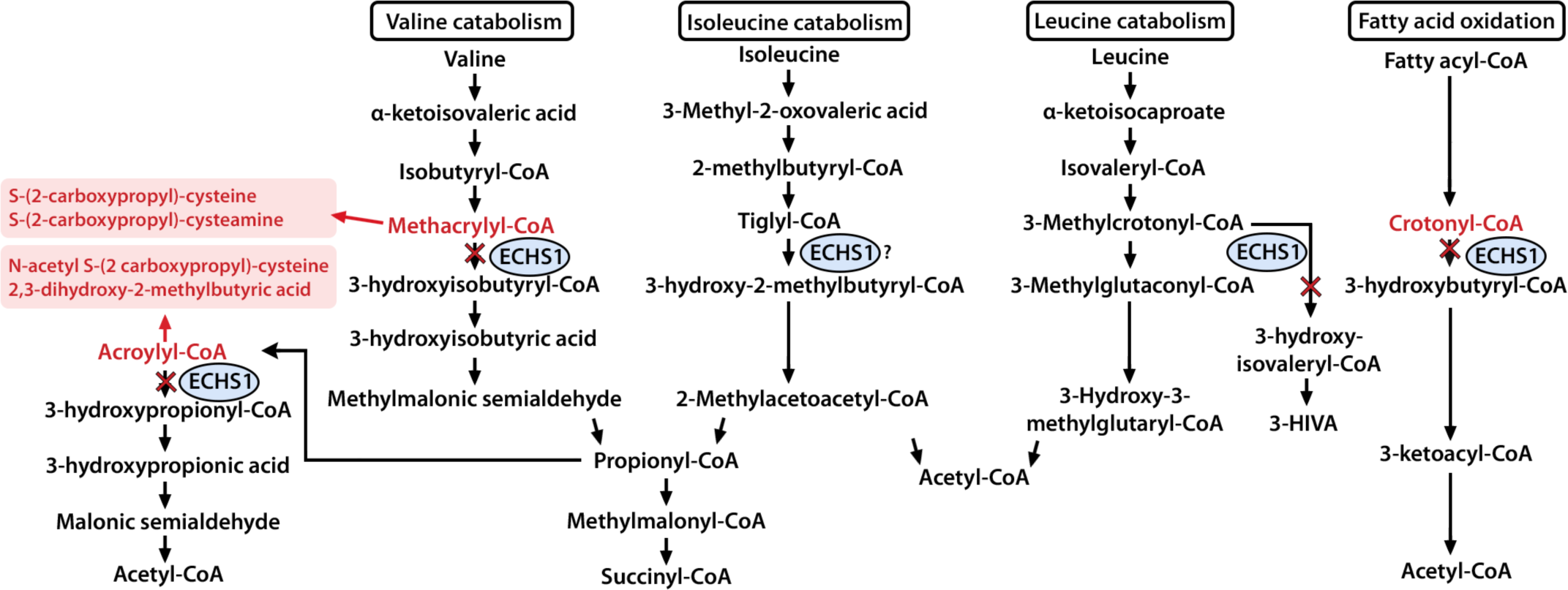
The roles of ECHS1 and abnormal metabolites formed in ECHS1D. Short-chain enoyl-CoA hydratase 1 (ECHS1) catalyses reactions in both fatty acid oxidation and valine, isoleucine, and leucine catabolism (branched-chain amino acids). Abnormal metabolites (red boxes) are all derived from the valine pathway. 3-HIVA = 3-hydroxyisovaleric acid.

There are currently no known effective therapeutic strategies for ECHS1D. Several diet and nutraceutical intervention trials on single patients have been reported (Table 1). Valine restriction^1,2,7,8^ or a low protein diet^5,9^ are reported to improve motor function in patients with slowly progressive or intermittent disease phenotypes, while more subtle improvements were observed in one patient with a more severe phenotype (Table 1). Other proposed therapies directed at restoring or bypassing the impairment in fatty acid catabolism, such as triheptanoin supplementation, have not been thoroughly assessed^10^. Overall, variability of treatment responses has hindered establishing consensus as to whether these treatments are beneficial or not. This highlights a need for a representative animal model system with which systematic treatment trials could be conducted.

**Table 1.**
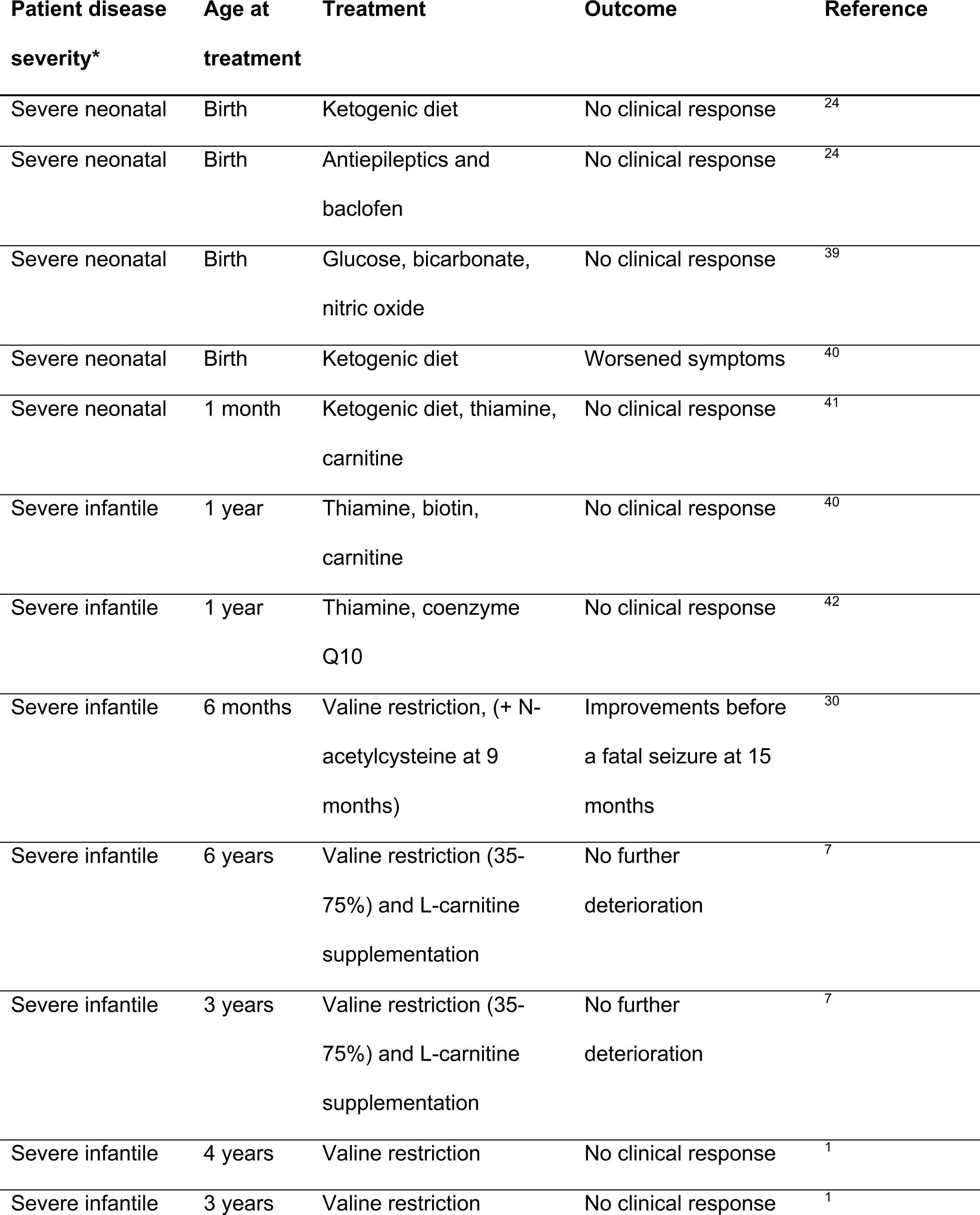
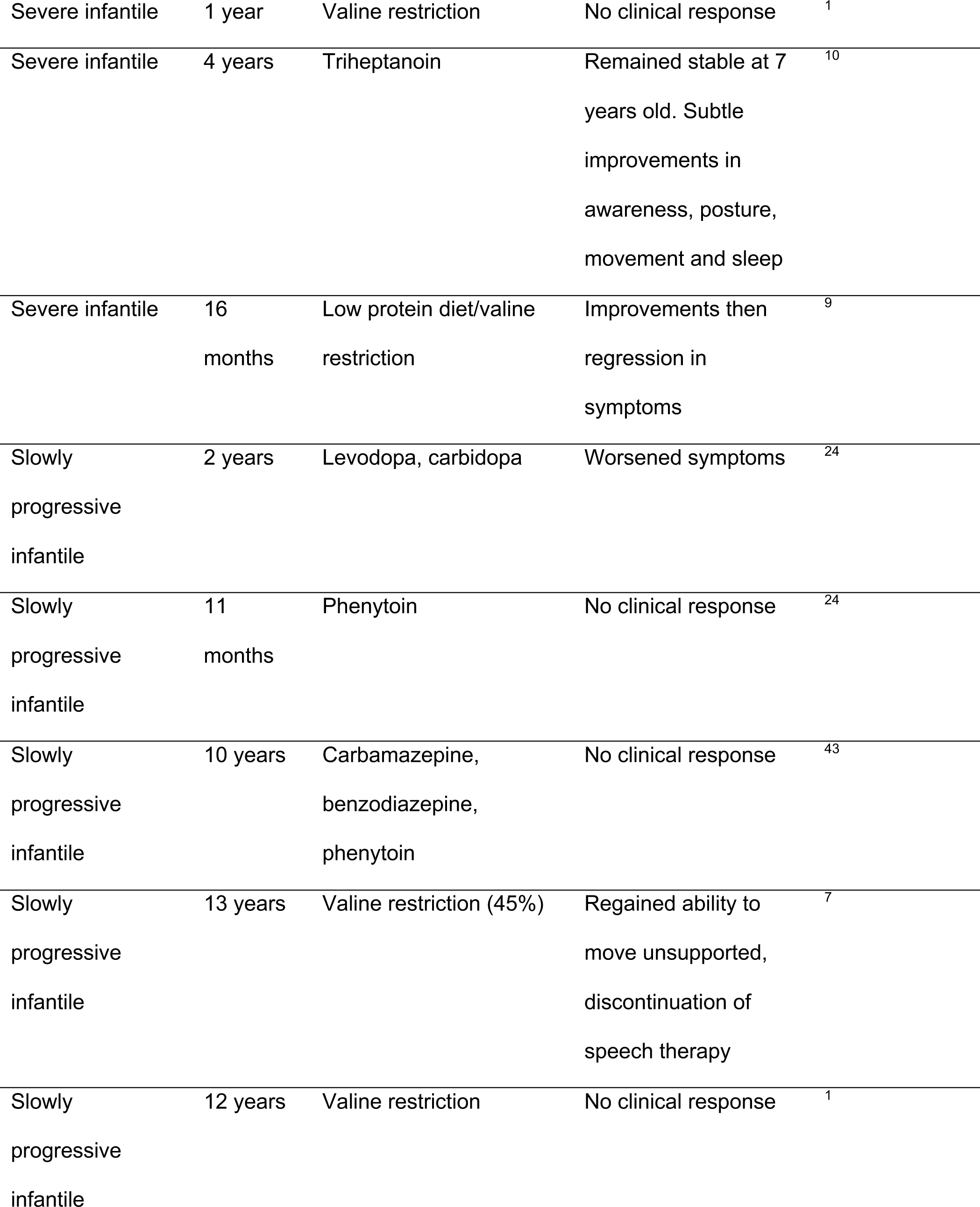
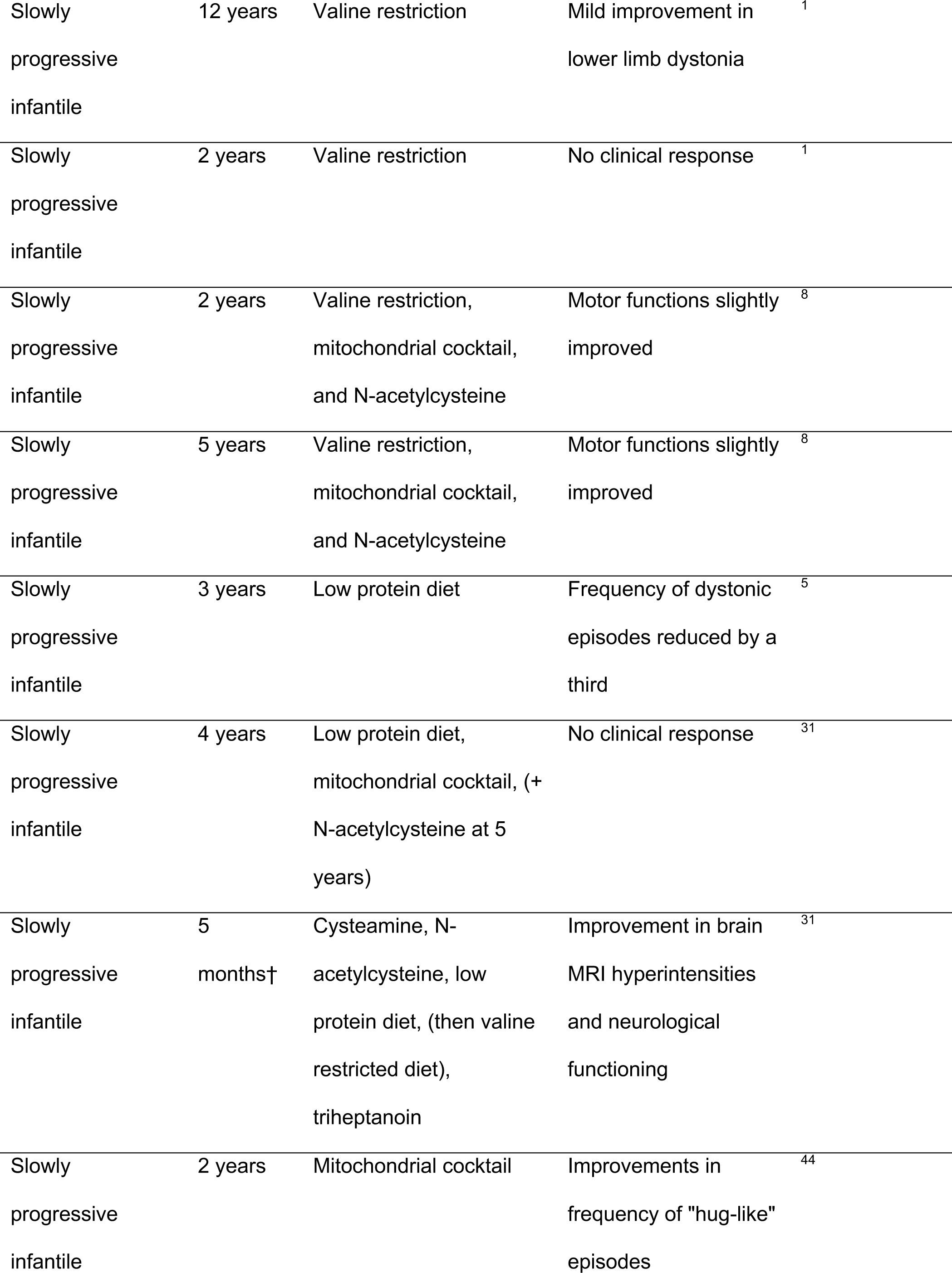
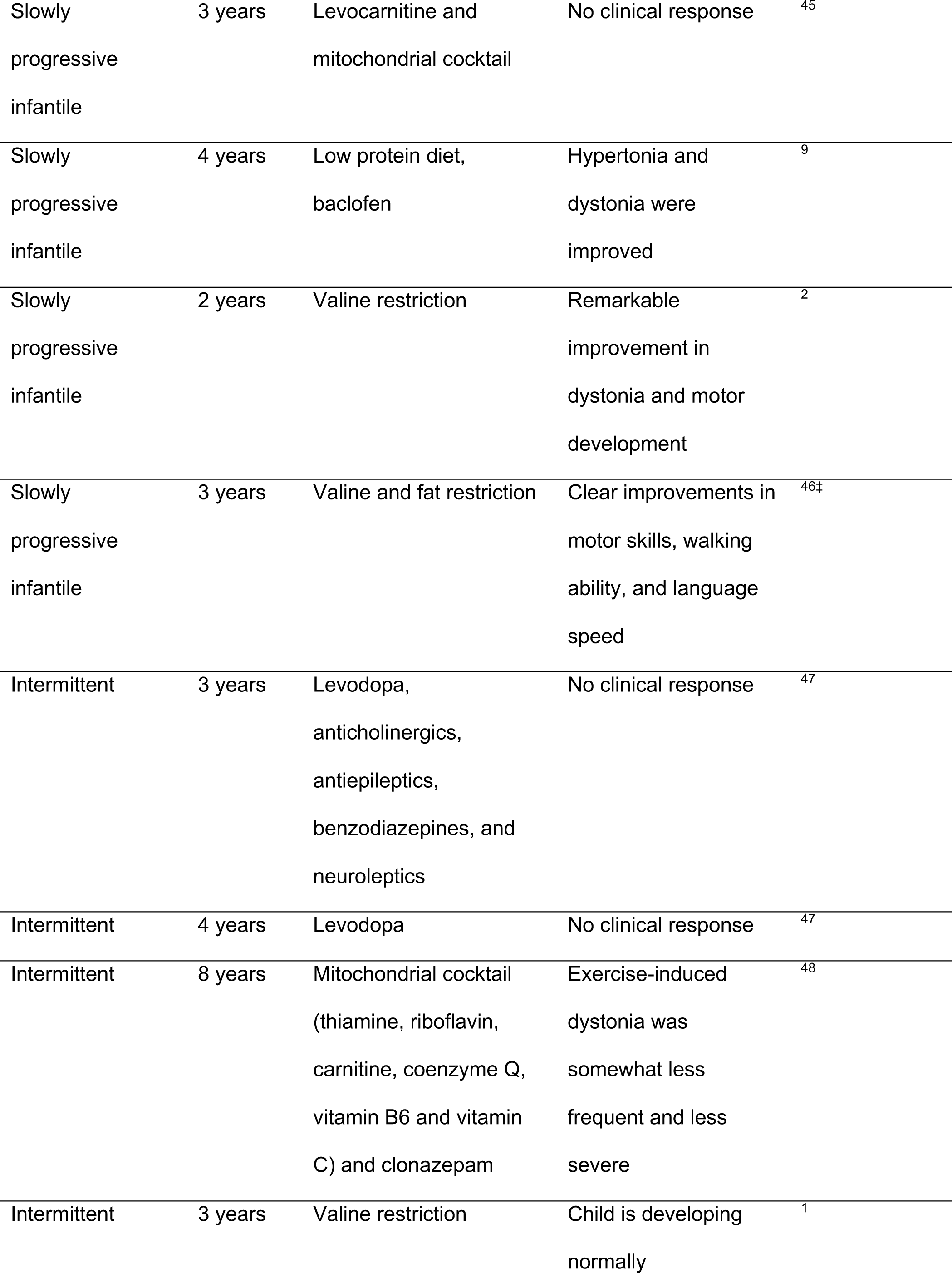
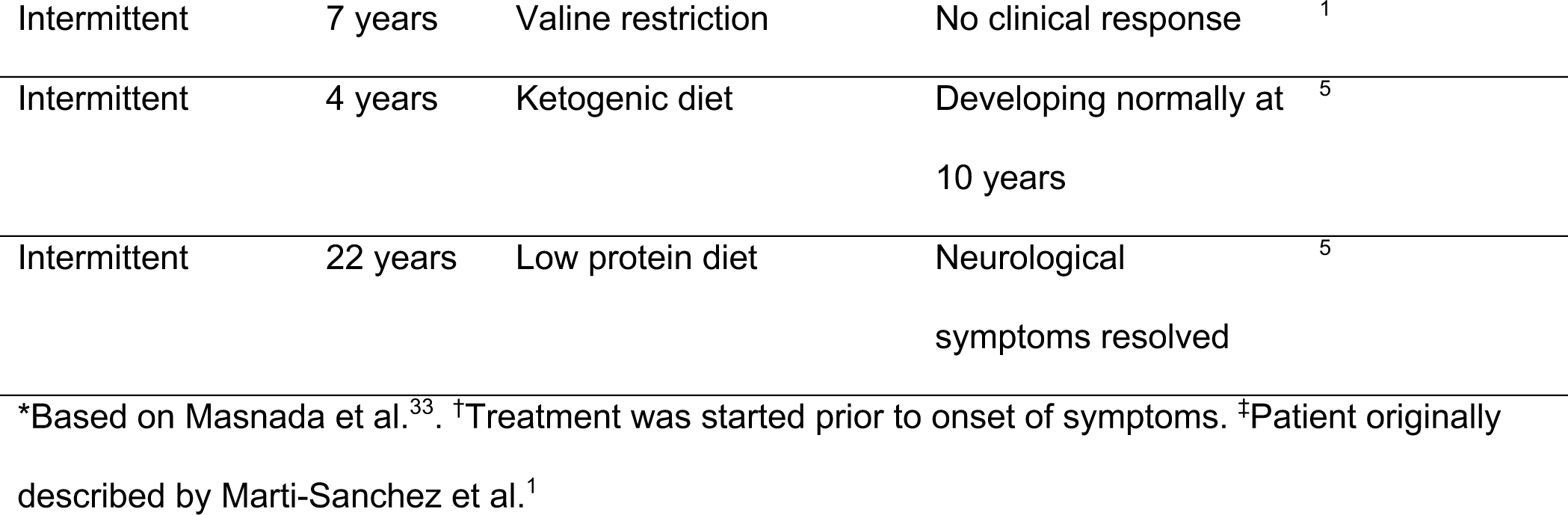
Treatments and outcomes in ECHS1 deficiency patients.

To date, no animal models of ECHS1D have been investigated for the purpose of treatment testing. The partial loss of ECHS1 has been investigated in mice and in human cells, frequently in the context of cancer proliferation due to the central role of ECHS1 in branched chain amino acid (BCAA) and fatty acid metabolism^11,12^. *Drosophila* have been extensively used to model human disease and have a synthetic customisable diet, making them a powerful tool for testing a range of dietary and nutraceutical interventions^13^. For this reason, we explored whether *Drosophila* could be used for animal modelling studies of ECHS1D. We identified the *Drosophila* gene *CG6543* (hereon referred to as *Echs1*) as a single highly conserved ortholog of human ECHS1. We then validated a *Drosophila* ECHS1-deficient strain and used this to investigate the physiological and metabolic effects of ECHS1D, as well as determined whether dietary modifications could improve fly health. These experiments revealed severe physiological impairments in the ECHS1D fly model and demonstrated diet-responses of clinical relevance. Our work establishes the first animal model of ECHS1D that recapitulates the critical characteristics of the disease and extends the use of *Drosophila* as a nutrigenomic model for treatment discovery.

## Materials and methods

### *Drosophila* stocks and maintenance

The following lines were obtained from the Bloomington Stock Centre: *w^1118^* (BL6305), y[1] w[*]; TI{GFP[3xP3.cLa]=CRIMIC.TG4.1}Echs1[CR00484-TG4.1] (BL79270, called *Echs1^TG4^* here), CyO-GFP balancer (BL9325), *w*[1118]; and *Df(2R)BSC383/CyO* (BL24407, called *Echs1^Df^* here). The *Echs1* allele was first outcrossed to *w^1118^* for removal of the yellow cuticle pigment marker y^1^ (to avoid impacts to tyrosine metabolism), then maintained over a fluorescently marked balancer chromosome (CyO-GFP). Experiments were conducted on *Echs1^TG4^* homozygous larvae. Fly stocks were maintained at 22°C on sugar-yeast food under natural photoperiod conditions and experiments were performed at 25°C.

### Generation of transgenic lines

UAS-human ECHS1 transgenic flies (hECHS1) were generated by gene synthesis of human *ECHS1* cDNA (LD24265, #7061) Clone ID OHu09094C from Accession NM_004092 (Genscript Inc). Human *ECHS1* has a single transcript. This clone was inserted into the plasmid pUASTattB using EcoRI and XbaI sites.^14^ Transgenics were made via phiC31 mediated integration into the landing site at 86Fb (WellGenetics, Taiwan).

### Synthetic diet preparation

Chemically defined media used in this study was prepared as described in Martelli et al.^15^ Briefly, sucrose, agar, low solubility amino acids (L-isoleucine, L-leucine, and L-tyrosine), and stock solutions of buffer, metal ions, and cholesterol were combined with MilliQ water using a magnetic stirrer. After autoclaving at 120⁰C for 15 minutes, the solution was cooled at room temperature to ∼70⁰C. Stock solutions for amino acids, vitamins, nucleosides, choline, inositol, and preservatives were added. Liquid food was dispensed into vials or 48-well plates. The unmodified synthetic diet is referred to as the Holidic diet.

### Dietary manipulations

The modifications to the Holidic diet include manipulation of amino acid levels (valine, isoleucine, leucine), L-carnitine hydrochloride (Merck, C0283-5G), and Glycerol trienanthate (triheptanoin, Sigma, 92655-5ML). Diets with varied amino acid concentrations were prepared by modifying the base Holidic diet recipe (Table S1). Carnitine was dissolved in Milli-Q water. Carnitine and triheptanoin were added to the Holidic diet prior to setting. To create starvation conditions, larvae were reared on 2% agar.

### Larva collection and sorting

To obtain large quantities of larvae that were developmentally synchronised, embryos were collected from population cages containing 80 females and 20 males allow to lay for 8 hours on apple juice agar plates supplemented with yeast paste. Embryos were collected and washed with distilled water in an embryo basket, then ∼50 µl of embryos were deposited onto plates containing the Holidic diet or modified diets as required. Plates were incubated for 24h at 25⁰C, then hatched larvae were transferred to the required diet mediums. To obtain homozygous mutant larvae, *ECHS1^TG4^/CyO-GFP* flies were self-crossed and selected on the absence of the fluorescent balancer. For controls, heterozygotes (*Echs1^TG4^/+*) or wildtype (*w^1118^)* was used. Heterozygotes were obtained by crossing *Echs1^TG4^/CyO-GFP* males to *w^1118^* females.

### Quantitative RT-PCR

Three replicates of 20 whole two-day old *Echs1^TG4^* homozygotes and *w^1118^* larvae were collected into microtubes containing 500 µl TRIzol (Bioline), homogenised and processed as directed by the manufacturer. RNA purity and concentration were evaluated by spectrophotometry (NanoDrop ND-1000, NanoDrop Technologies). cDNA synthesised using oligoDT and random hexamers (Bioline). Quantitative PCR was performed on a lightcycler (Thermo Fisher) with SYBR chemistry. The following primer pairs were designed using NCBI Primer BLAST: *Echs1*: F 5’-TTA AGG AGA TGG TGG GCA AC-3’; R 5’-ACC AAA CTT GGC CTT GTC AC-3’, *RpL11*: F 5’-CGA TCC CTC CAT CGG TAT CT-3’; R 5’-AAC CAC TTC ATG GCA TCC TC-3’. Reactions were run in triplicate and fold changes determined using the ΔΔCT method and the ribosomal protein L11 gene (*RpL11*) was used as a reference for normalising expression levels. The results were analysed by the Shapiro-Wilk normality test (p>0.05), before comparing the transcript levels between the two treatment groups using two-tailed Student’s t-tests (p < 0.05) (GraphPad Prism 9).

### Developmental timing and viability

To measure viability and time of death of *Echs1^TG4^* larvae, newly hatched larvae were transferred into 48-well plates with one larva per well. Wells contained 700 µl of the Holidic diet or modified treatment diet. Plates were covered with clear PCR plate film with 20 small ventilation holes per well made using a 20-gauge needle. Forty-eight wells per genotype per treatment were imaged once an hour for 14 days using a custom-made robotic imaging system with infrared illumination at 25°C in darkness (T.K. Johnson, unpublished). Image series from each well were collated in ImageJ and assessed for the hour at which each larva stopped moving (time of death). Hour of death was compared between treatment and genotype groups using a two-way ANOVA with multiple comparisons and Tukey’s HSD. Error bars represent SEM unless specified.

### Larval phenotypic assessments

Larval crawling ability was assayed using a described method.^16^ Briefly, two-day old *Echs1^TG4^* larvae were collected by adding a 20% w/v sucrose solution to population vials, the food was gently disrupted using a small paintbrush to release and float the larvae. This mix was poured through a fine cloth mesh to isolate the larvae from the food. Larvae were then rinsed with PBS. Five larvae per genotype per treatment for five replicates were collected. Larvae were transferred to 35 mm petri dishes containing 2% apple juice agar dyed blue (blue food dye, Queen brand) and allowed to acclimate for 3 minutes. Larvae were then recorded for 3 minutes with a webcam (Logitech). Movies were converted to .avi format using ffmpeg (https://ffmpeg.org), then processed and analysed in ImageJ using the wrMTrck plugin (https://www.phage.dk/plugins/wrmtrck.html). For each larva, the total length of movement (mm) and average speed (mm/s) were calculated. To measure growth rate of *Echs1^TG4^*, embryos were collected and deposited on 6 plates containing sugar-yeast food. Plates were incubated at 25°C and every 24 hours a plate was retrieved, and individual larvae were transferred to a petri dish containing 2% agar dyed blue for contrast between the larva and the background. To keep larvae still the plate was pre-cooled on ice. Images were taken using a Leica AF6000 LX microscope camera and size was obtained by measuring the area of larvae using the square tool in ImageJ.

### Metabolomic profiling

Five replicates of 30 two-day old *Echs1^TG4^* larvae were collected, washed in PBS, blotted dry, and weighed. Samples were transferred to 1.5 mL safe lock microtubes (Eppendorf), flash frozen in liquid nitrogen, and stored at -80°C. Larvae were thawed and then homogenised using a disposable pestle in 20 µL of ice-cold extraction solvent consisting of a 2:6:1 chloroform:methanol:water ratio with 2 µM of (CHAPS, CAPS, PIPES and TRIS) acting as internal standards. Once homogenised, additional solvent was added to a final ratio of 20 µL extraction solvent per mg of larvae. Samples were vortexed for 30 seconds and sonicated in an ice-water bath for 10 minutes, then centrifuged at 4°C (22,000 ×g for 10 min). The supernatant was transferred to a glass vial for LC-MS metabolic analysis. Twenty microlitres of each extract was combined to make a pooled quality control sample. Heterozygotes were used as a control.

LC–MS was performed using a Dionex Ultimate 3000 UHPLC coupled to a QExactive Plus mass spectrometer (Thermo Scientific). Samples were analysed by hydrophilic interaction liquid chromatography (HILIC) following a previously published method.^17^ The chromatography utilised a ZIC-p(HILIC) column 5µm 150 x 4.6 mm with a 20 x 2.1 mm ZIC-pHILIC guard column (both Merck Millipore, Australia) (25 °C). A gradient elution of 20 mM ammonium carbonate (A) and acetonitrile (B) (linear gradient time-%B: 0 min-80%, 15 min-50%, 18 min-5%, 21 min-5%, 24 min-80%, 32 min-80%) was utilized. Flow rate was maintained at 300 μL/min. Samples were stored in the autosampler (6°C) and 10 μL was injected for analysis. Mass spectrometry was performed at 70,000 resolution operating in rapid switching positive (4 kV) and negative (−3.5 kV) mode electrospray ionization (capillary temperature 300°C; sheath gas flow rate 50; auxiliary gas flow rate 20; sweep gas 2; probe temp 120°C). Samples were randomized and processed in a single batch with intermittent analysis of pooled quality-control samples to ensure reproducibility and minimize variation. For accurate metabolite identification, a standard library of ∼500 metabolites were analyzed before sample testing and accurate retention time for each standard was recorded. This standard library also forms the basis of a retention time prediction model used to provide putative identification of metabolites not contained within the standard library.

### Metabolic profiling data analysis

Acquired LC-MS/MS data was processed in an untargeted fashion using open-source software IDEOM, which initially used msConvert^18^ (ProteoWizard) to convert raw LC-MS files to mzXML format and XCMS to pick peaks to convert to .peakML files. Mzmatch was subsequently used for sample alignment and filtering.^19,20^ Metabolites were identified based on accurate mass (<2 ppm) and comparison of their retention time against that determined for compounds in the standard library or predicted on the basis of their physiochemical characteristics^20^. Only metabolites that were identified with a level of confidence equal to or greater than 6 in IDEOM were used for downstream functional and statistical analyses, using MetaboAnalyst 6.0.^21^ Global metabolic variations due to genotype and diet were visualized using PCA, to verify the contribution of each metabolite in the variance of each treatment. Two-sample t-tests [false discovery rates (FDR) < 0.05] and fold change analysis (2.0 cutoff) was used to identify significant changes in metabolite levels between mutant and control.

## Results

### ECHS1 deficiency in *Drosophila* causes severe survival, growth, and behavioural defects

We first identified *Drosophila CG6543* (hereon referred to as *Echs1*) as the sole ortholog of human *ECHS1* with 71% sequence similarity and a conserved enoyl-CoA hydratase domain.^22^ To assess the consequences of loss of *Echs1* in *Drosophila*, we acquired a putative null allele of *Echs1* (referred herein as *Echs1^TG4^*) comprising an artificial disruptive exon inserted into the first intron using CRISPR/Cas9 technology.^23^ The cassette contains a splice acceptor followed by a T2A peptide cleavage sequence, the GAL4-coding sequence, and a polyA signal sequence (Figure 2A). This is expected to severely truncate *Echs1* gene products and yield expression of the transcriptional activator GAL4 in the same spatial and temporal pattern as *Echs1*. Consistent with this, quantitative RT-PCR showed *Echs1* transcripts were absent in *Echs1^TG4^* homozygotes compared to wild-type controls (Figure 2B). This allele therefore causes complete loss of *Echs1*, representing the most severe form of ECHS1D, much like the truncating p.(Ala31Glufs*23) patient variant.^4^

**Figure 2.**
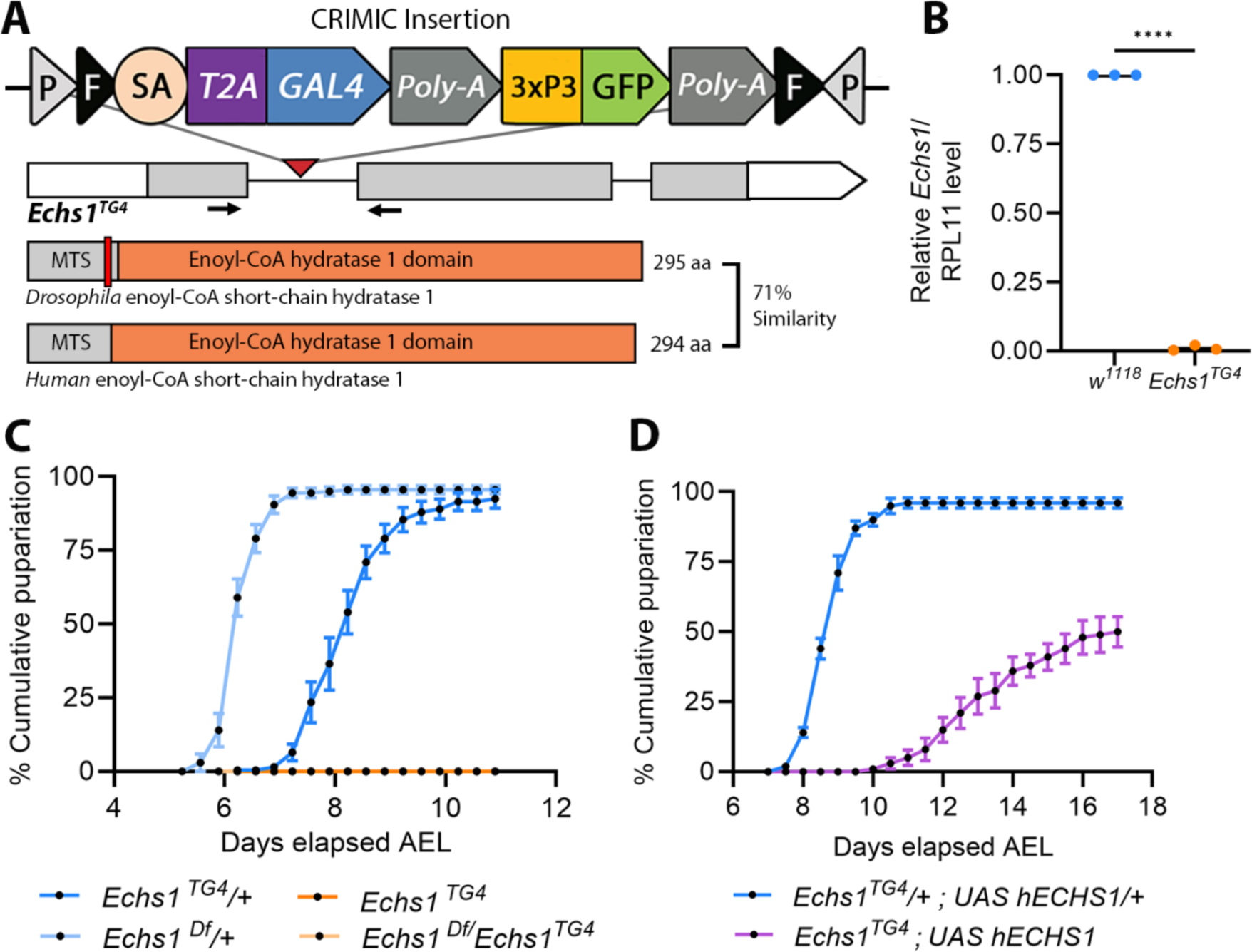
Loss-of-function mutations in *Echs1* in *Drosophila* cause lethality. (A) Schematic of the *Echs1^TG4^* allele in *Drosophila* containing the disruption T2A-GAL4 (TG4) cassette. Disruption causes a truncation within the predicted mitochondrial targeting sequence (MTS). (B) Quantitative RT-PCR of *Echs1* using homogenates of 2^nd^ instar *Echs1^TG4^* homozygote larvae and age-matched *w^1118^* (control) showing no *Echs1* expression. (C) Cumulative pupariation rates of *Echs1^TG4^*, *Echs1^TG4^*/*Echs1^Df^* (complementation), and heterozygote controls. Five replicate vials tested per genotype, 20 individuals per replicate. (D) Rescue of larval-lethal phenotype with UAS-human ECHS1 (*hECHS1*) expressed in a *Echs1^TG4^* background. AEL = after egg lay. Students t-test: ****p< 0.0001, ***p<0.001.

A characterisation of *Echs1^TG4^* revealed that *Echs1* loss in *Drosophila* is lethal during the larval stage as individuals were unable to reach pupariation (Figure 2C). It was confirmed that this phenotype was not due to other recessive lethal mutations on the *Echs1^TG4^* chromosome by complementation using an independently derived deletion allele spanning *Echs1* (*Echs1^Df^*). Compound heterozygotes (*Echs1^TG4^*/*Echs1^Df^*) similarly failed to pupariate (Figure 2C). The pre-pupal arrest of *Echs1^TG4^* was partially rescued by the expression of a human *ECHS1* transgene (UAS-*hECHS1*) driven by GAL4 produced by the *Echs1^TG4^* locus (Figure 2D).

Closer analysis of the larval period revealed that 50% of *Echs1^TG4^*homozygotes and compound heterozygotes (*Echs1^TG4^*/*Echs1^Df^*) died by day 6 post-egg-lay as second instar-staged larvae (Figure 3A). Consistent with this, *Echs1^TG4^* fail to grow beyond 48h post egg-lay and typically arrest at this size prior to 9 days post-egg-lay (Figure 3B). A locomotor activity assay was performed to assess neuromuscular ability of larvae at 48h (before the effects of growth impairment). *Echs1^TG4^* larvae crawled significantly shorter distances and at a slower average speed than controls (Figure 3C-D). Together, these phenotypes resemble a severe course of ECHS1D observed in humans consistent with the nature of the mutation.

**Figure 3.**
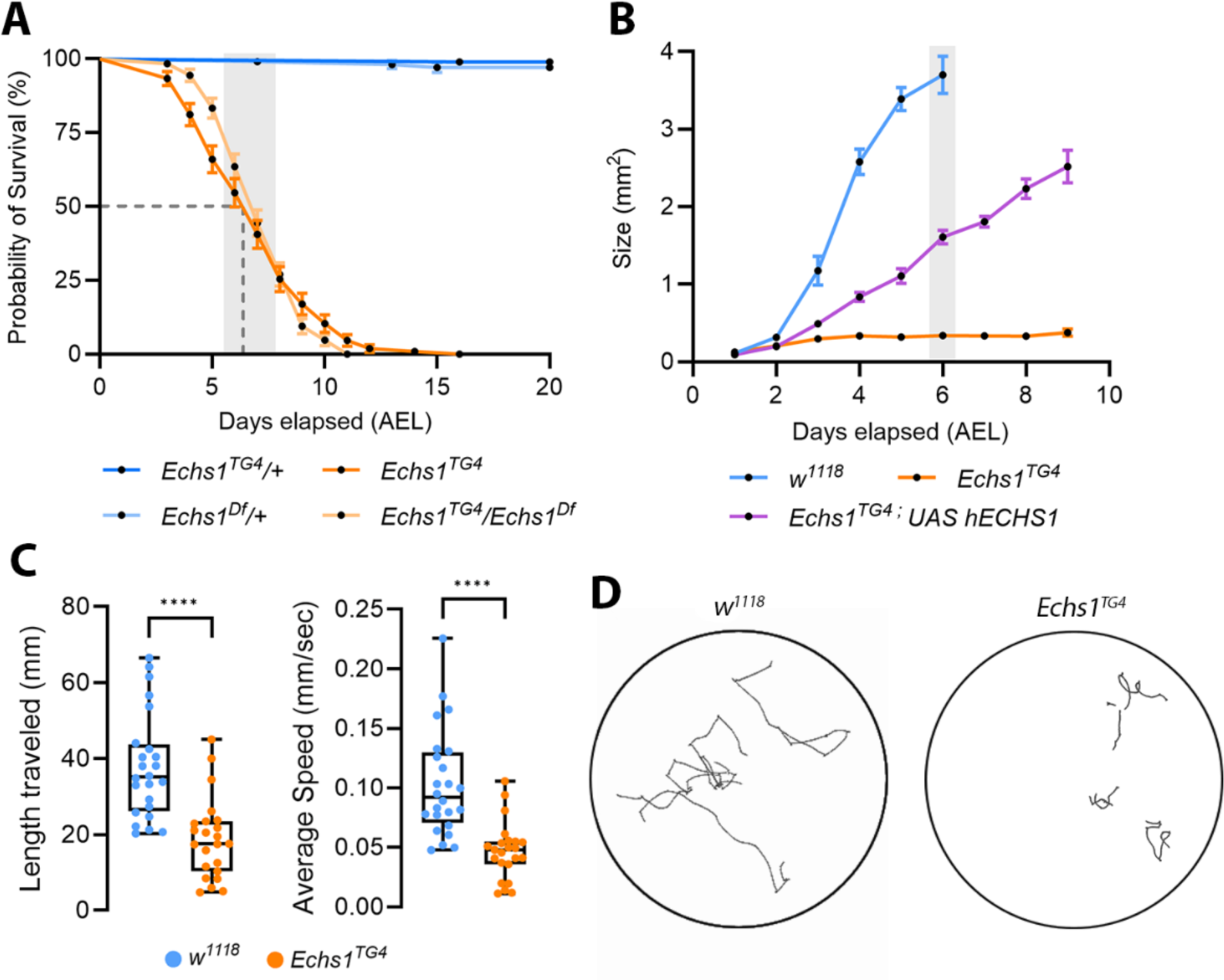
Loss of *Echs1* causes larval arrest and impairs growth and locomotor activity. (A) Survival curves of *Echs1^TG4^, Echs1^TG4^/Echs1^Df^* and controls reared on a sugar-yeast diet. Five replicate vials were tested per genotype with 20 individuals per replicate. Grey rectangle indicates the time-window when control larvae progressed to pupariation. (B) Larval growth curves of *Echs1^TG4^*, *Echs1* human rescue (*Echs1^TG4^;UAS hECHS1*) larvae, and controls. 20 larvae measured per time point per genotype. Grey rectangle indicates the window of pupariation time for control (*w^1118^*). (C) Locomotion and (D) larval movement tracks of 2^nd^ instar *Echs1^TG4^* homozygotes. Five larvae were tested per replicate, with five replicates tested per genotype. T-test: ****P < 0.001. AEL = After egg lay.

### Loss of *Echs1* causes broad changes to metabolism

We next wanted to explore how loss of *Echs1* impacts fly metabolism, both with respect to the profile observed in ECHS1D patients, and to shed light on disease processes. For this we performed untargeted metabolomics on *Echs1^TG4^* and heterozygous controls (*Echs1^TG4^/+*) sampled at 48h old to capture early changes in disease progression. Principal component analysis on the 1,305 putatively identified metabolites clearly distinguished the ECHS1D state from that of the control (Figure 4A). Overall, we detected 379 significantly affected metabolites in the *Echs1^TG4^* larvae. We first looked for the accumulation of valine-derived metabolites proposed to be the pathogenic cause of ECHS1D^24^ (Figure 1). Most notably, we found S-(2-carboxypropyl)cysteamine and 2,3-dihydroxy-2-methylbutyric acid, derived from the ECHS1 substrates methacrylyl-CoA and acryloyl-CoA respectively, both elevated by 6-fold in *Echs1^TG4^*, consistent with patient data (Figure 4B). We also observed other elevated metabolites in the fly model reported to be similarly altered in ECHS1D patients such as phenyllactic acid (8-fold), 2-methylcitric acid (7-fold), and 3-hydroxyisovaleric acid (5-fold) (Figure 4B-C). These data provide strong support that our ECHS1D model resembles the human disorder at the metabolic level. Other known elevated metabolites such as lactic acid, alanine, and short-chain acylcarnitines were unchanged or reduced in the model (Figure S2). Key metabolites of energy metabolism were assessed, as oxidative phosphorylation enzymes have been shown to have reduced activity in some severe cases of ECHS1D^25,26^, however these metabolites were unchanged in the fly model (Figure S2).

**Figure 4.**
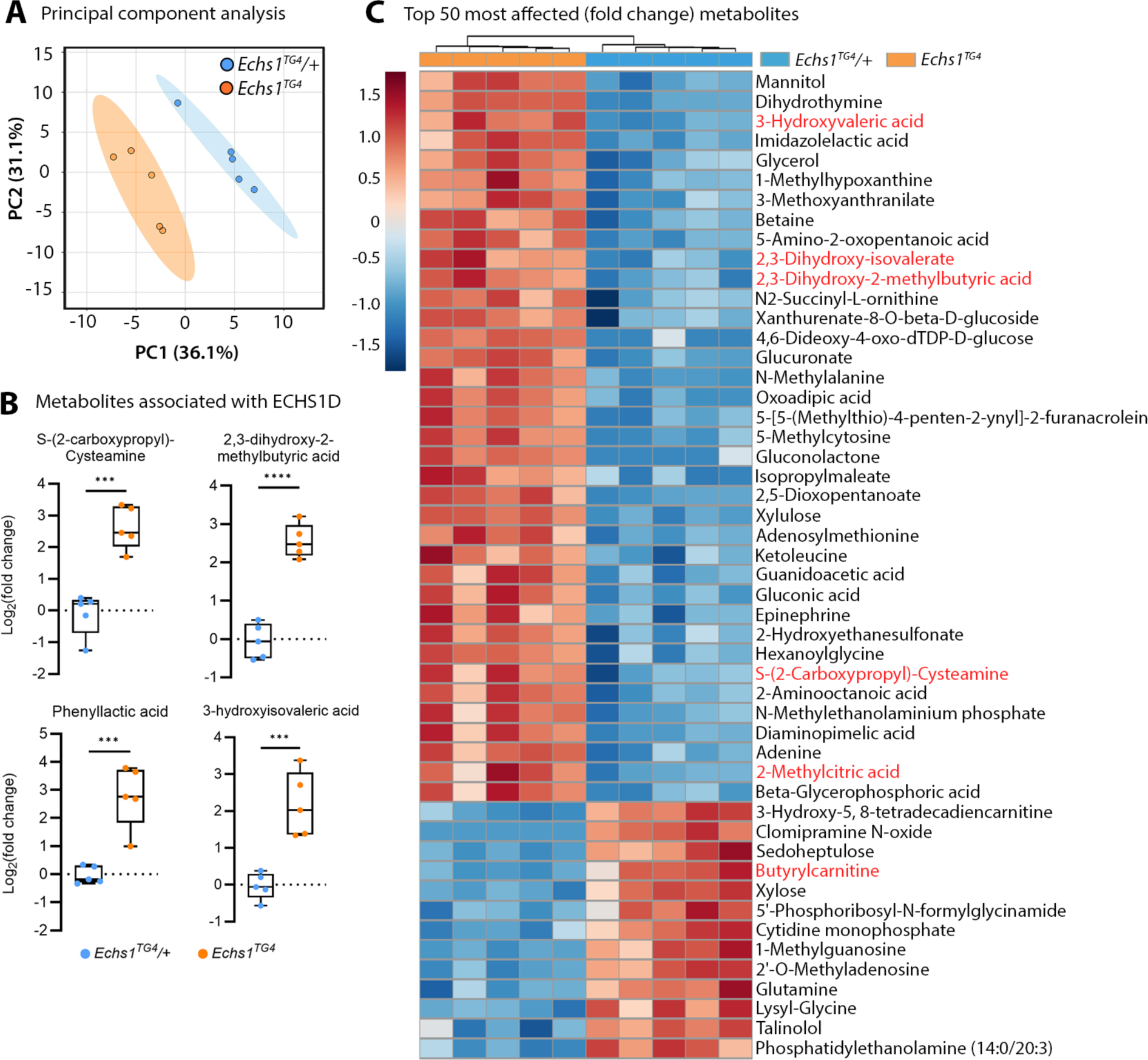
Metabolic profile of *Echs1*-deficient *Drosophila* larvae. (A) Principal component analysis (PCA) of 1,305 putative metabolites: circles indicate 5 biological replicates of 30 homogenised whole larvae, shaded ellipses represent 95% confidence intervals. (B) ECHS1 deficiency-associated metabolite levels (Log2(fold change)) compared to control *Echs1^TG4^*/+ larvae. (C) Heatmap of the top 50 most-affected metabolites between *Echs1^TG4^* and *Echs1^TG4^/+* (T-test P < 0.05, FDR). Only metabolites with a human metabolome database (HMDB) reference are included. Metabolites in red have been associated with human ECHS1D.

Next, we looked beyond BCAA metabolism and known markers ECHS1D pathogenesis. The most highly elevated metabolites were associated with carbohydrate metabolism (Figure 4C, Figure S1A-B). These included gluconolactone (30-fold), glucuronate (21-fold), and gluconic acid (15-fold). Gluconolactone and gluconic acid are derived from the oxidation of glucose in insects via glucose dehydrogenase.^27^ Glucuronate, also derived from glucose, functions in xenobiotic detoxification and can be proinflammatory.^28,29^ This therefore may reflect an effort to protect against cellular damage or a state of chronic inflammation. An enrichment analysis identified glutathione-associated metabolites are elevated in *Echs1^TG4^*larvae consistent with the mounting of a protective response. We also noted elevated pentose-phosphate pathway metabolites which could suggest a need to replenish NADPH, possibly to support the antioxidant activity of glutathione.

### Dietary valine restriction prolongs the survival of *Echs1*-deficient flies

Dietary restriction of valine has been trialled widely in ECHS1D patients, but outcomes have been variable.^1,2,5,7–9,30,31^ Before we could begin testing the ECHS1D fly model with dietary valine restriction we first wanted to confirm that *Echs1^TG4^*larvae were feeding and not simply arresting due to starvation. We found that starvation resulted in significantly shorter time to death for larvae compared to fed conditions, and on par with controls, suggesting *Echs1^TG4^*larvae were indeed consuming the diets provided (Figure 5A). When we raised larvae on synthetic diets with reduced valine content, we observed significant extensions to survival (for 0, 25, and 50% of the full diet complement, Figure 5B). Reduction of valine to 25%, for example, extended survival to a mean of 8.3 days compared to 5.6 days at 100% valine (Figure 5B). Control flies, however, displayed a dose-dependent reduction in pupariation rate with the same valine concentrations (0, 25, and 50% valine), all negatively impacting developmental progression (Figure 5C). These data suggest that, at least in *Drosophila*, there is a delicate balance between the positive and negative impacts of valine restriction in ECHS1D.

**Figure 5.**
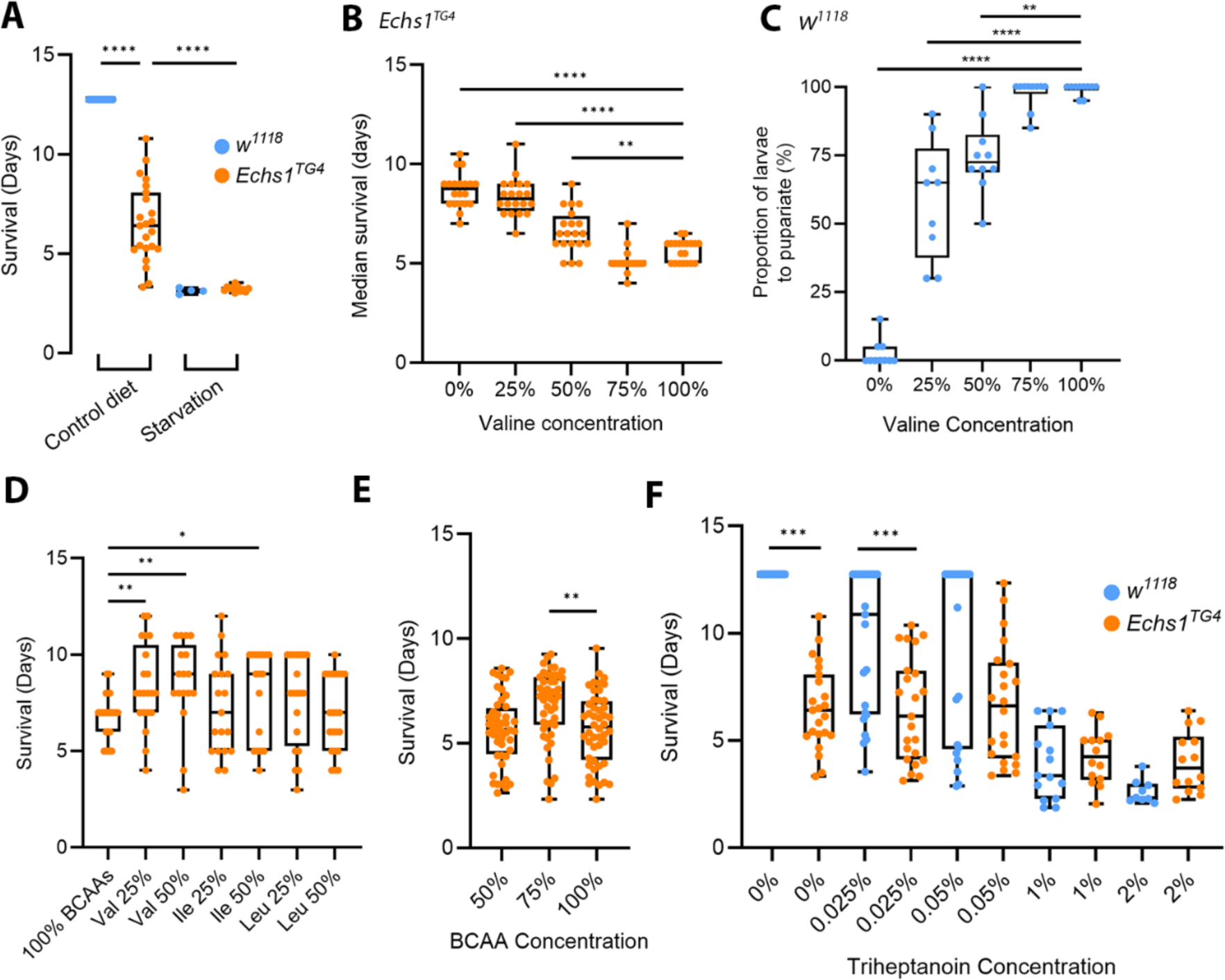
*Echs1^TG4^* larvae respond to valine, isoleucine, and total BCAA restriction. (A) Larval survival of *Echs1^TG4^* under starvation conditions (agar only). (B) Median larval survival of *Echs1^TG4^*on valine-restricted diets. Percent dietary valine values are relative to the complete diet. Twenty larvae were tested per replicate, with 10 replicates per treatment. (C) Proportion of *w^1118^* larvae that pupariated on valine-restricted diets. Twenty larvae were tested per replicate, with 5 replicates per treatment. (D) Larval survival on diets containing dilutions of each branched-chain amino acid (Val = valine, Ile = isoleucine, and Leu = leucine). (E) Larval survival on diets depleted of branched-chain amino acids. Percentages are relative to the complete diet. (F) Larval survival on diets supplemented with triheptanoin (7-carbon fatty acid). Students t-test: ****P < 0.0001, ***P < 0.001, **P < 0.01, * P < 0.05

To test whether the survival extension was valine-specific we also tested restriction of the other BCAAs. Isoleucine restriction (to 50%, but not 25%) also significantly extended *Echs1^TG4^* larval survival, whereas leucine restriction had no effect (Figure 5D). This suggests that in *Drosophila*, Echs1 and the pathogenesis of ECHS1D may involve both valine and isoleucine metabolism.

We have previously observed that restricting a single BCAA can cause developmental delay due to a BCAA imbalance rather than a deficiency.^15^ To test whether amino acid imbalance was restricting developmental progression in *Echs1^TG4^* larvae, all BCAAs were restricted equally (to 50 and 75% relative to the complete diet). At 75%, but not 50% BCAA concentration, we observed a significant survival extension equivalent to reducing valine alone to 50% (Figure 5E). Notably, at this BCAA content, we do not observe negative effects on developmental progression or survival on control flies.^15^ This suggests that the extension in *Echs1^TG4^*survival is likely due to reduced BCAA pathway flux, while their inability to progress further in development is not likely due to a BCAA deficiency.

This observation made us curious as to whether other roles of ECHS1 might be contributing to the severe phenotype in flies, and perhaps in humans. ECHS1 is also required for the catabolism of short-chain fatty acids (Figure 1), but the contribution of fatty acid β-oxidation to the ECHS1D phenotype remains unclear. It has been proposed that the metabolic blockage in fatty acid β-oxidation could be bypassed by supplementing medium odd-chain fatty acids (such as the triglyceride triheptanoin), which are metabolised via an alternate pathway thereby providing anaplerotic substrates.^32^ Administering varying dosages of triheptanoin did not extend survival of *Echs1^TG4^*compared to the control diet, and control larvae had a marked decrease in survival as dosage increased (Figure 5F). This suggests that the products of fatty acid metabolism are not a limiting factor in ECHS1D.

## Discussion

ECHS1D is a severe childhood disorder that lacks effective treatment options and a model system in which to test the efficacy of proposed interventions. Here we established *Drosophila* as a model for investigating ECHS1D pathogenesis and assessing the effectiveness of dietary modification as treatment for this inherited metabolic disorder. We found that loss of *Echs1* in *Drosophila* causes poor motor behaviour, growth stagnation, and premature mortality; closely resembling a severe course of ECHS1D^33^. Truncating mutations in *ECHS1* have been observed in humans, and much like our *Drosophila Echs1^TG4^* allele, these are associated with a rapidly fatal condition.^4^ We also found metabolites that are routinely identified in ECHS1D patients and linked to ECHS1D pathogenesis, to be highly elevated in the *Drosophila* model.^6^ Thus, the evidence presented here, in addition to the partial rescue of *Echs1^TG4^* survival upon human *ECHS1* expression, demonstrate strong evolutionary conservation between *Drosophila* and humans in both ECHS1-related metabolic systems and disease processes. This model therefore could be utilised in drug and nutrient screens to further understand disease mechanisms and develop treatments for ECHS1D.

Clinical reports have indicated that valine-restricted diets may prevent or lessen ECHS1D symptoms, however outcomes have been variable.^30,31^ Here, we were able to restrict valine intake specifically and for the entire life of *Echs1^TG4^*larvae, and this prolonged survival in a dose-dependent manner, suggesting benefit from the dietary intervention. However, since valine is an essential dietary component, restriction can cause deleterious effects on fly health compared to restriction of equal amounts of all BCAAs.^15^ This may be the reason why health restoration was limited to a modest extension of mean survival. Together these data suggest either that there are other pathogenic mechanisms in ECHS1D operating beyond the valine pathway, or that dietary restriction of valine alone is unable to reduce pathway flux sufficiently to avoid metabolic intoxication.

It is currently unclear whether other pathways are impacted in ECHS1D.^25,26^ Here, we observed elevated metabolites in the leucine pathway suggesting it may also be impacted. One of these is 3-hydroxyisovaleric acid (3-HIVA) observed as elevated in both the fly model and in ECHS1D patients.^34,35^ Elevated 3-HIVA is typically suggestive of a perturbed leucine metabolism, arising either by conversion from the keto-acid of leucine, α-ketoisocaproate (α KIC) by α-KIC dioxygenase, or from elevated methylcrotonyl-CoA which is processed to 3-HIVA by ECHS1^36,37^ (Figure 1). Given that here we observed 3-HIVA in the context of *Echs1* deficiency, we infer that α-KIC is the likely source of 3-HIVA, potentially suggesting reduced activity of the upstream branched chain ketoacid dehydrogenase complex. Interestingly, α-KIC dioxygenase recognises 4-hydroxyphenylpyruvate and phenylpyruvate (from tyrosine and phenylalanine metabolism, respectively) as substrates.^38^ Both of these metabolites, along with their lactate derivatives 4-hydroxyphenyllactate and phenyllactate (also elevated in patients) were highly elevated in *Echs1* deficient larvae. Whether there is dysregulation α-KIC dioxygenase in the context of ECHS1D and this connection between BCAA and tyrosine/phenylalanine metabolism, warrants further investigation.

The contribution of the fatty acid oxidation pathway to ECHS1D is also unclear. Here, triheptanoin feeding did not improve survival of the model, suggesting that the products of fatty acid oxidation (propionyl-CoA and acetyl-CoA) are not in short supply in *Drosophila* larvae with ECHS1D. Consistent with this idea, acetyl-CoA was unchanged in the metabolic profile. It is possible that the ECHS1 protein has roles beyond both valine and fatty acid metabolism, potentially involving maintenance of oxidative phosphorylation enzymes.^26^ Genetic interaction studies may prove useful for determining the extent to which this is the case. Given that *Drosophila* are highly amenable to these experiments, we anticipate that this model will be valuable not only as a screening tool for treatment discovery, but also for exploring the complex mechanisms that underpin ECHS1D pathogenesis.

## Limitations of the study

There are physiological differences between model organisms, such as *Drosophila* used here, and humans which could manifest as differences in both biochemical profiles and treatment responses. A caveat of the untargeted metabolic analysis is some metabolites were annotated based on the accurate measurement of their mass which enables assignment at a molecular formula level. Further support for their identification by comparison against authentic standards is needed. Additionally, the treatments provided to our flies were administered prior to disease onset and for their entire lifespan. This does not reflect the typical treatment course in humans with ECHS1D, where specialty valine depleted diets are provided post-diagnosis.

## Data availability

Data that support the findings of this study are available from the corresponding authors upon request.

## Acknowledgements

We thank the Bloomington Drosophila Stock Centre, and Professor Hugo Bellen and Dr Oguz Kanca (Baylor College of Medicine) for fly stocks, the Australian Drosophila Biomedical Research Facility (Ozdros) for stock importation, and members of the Johnson and Piper labs for helpful discussions.

## Author contributions

Conceptualization, M.D.W.P. and T.K.J.; methodology, S.M.; investigation, S.M., F.M., C.K.B., and G.J.; visualization and analysis, S.M., F.M., C.K.B., R.B.S., M.D.W.P., and T.K.J.; funding acquisition, M.D.W.P. and T.K.J.; project administration, M.D.W.P. and T.K.J.; supervision, J.C., S.D., M.D.W.P., and T.K.J.; writing, S.M., M.D.W.P., and T.K.J

## Supplementary data

### Figures

**Figure S1.**
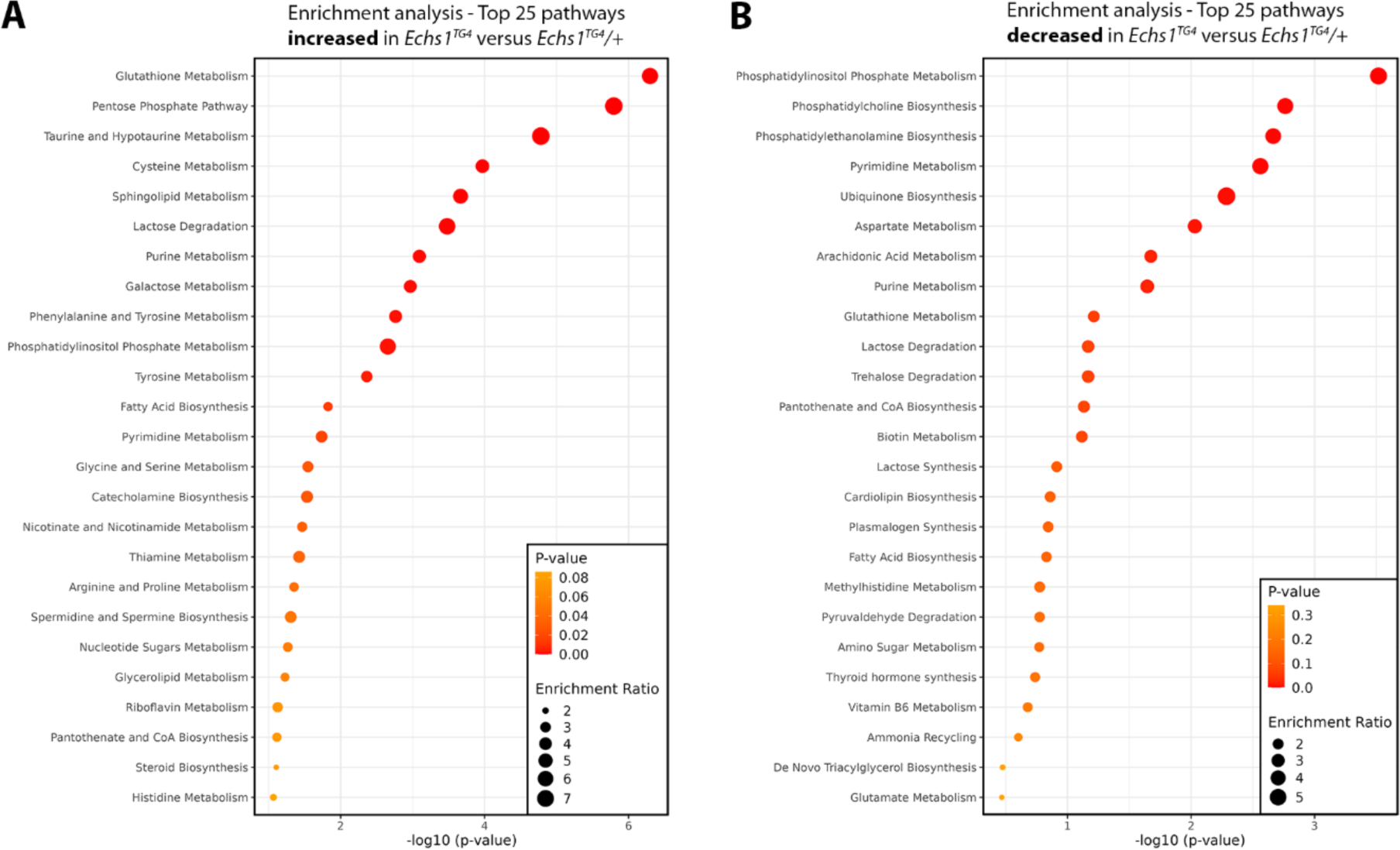
Metabolome enrichment analysis. (A, B) Quantitative enrichment analysis showing the top 25 metabolic pathways (A) increased or (B) decreased in *Echs1^TG4^* versus *Echs1^TG4^*/+ reared on the Holidic diet. Enrichment analysis based on the small molecule pathway database (SMPDB), performed in MetaboAnalyst 6.0.

**Figure S2.**
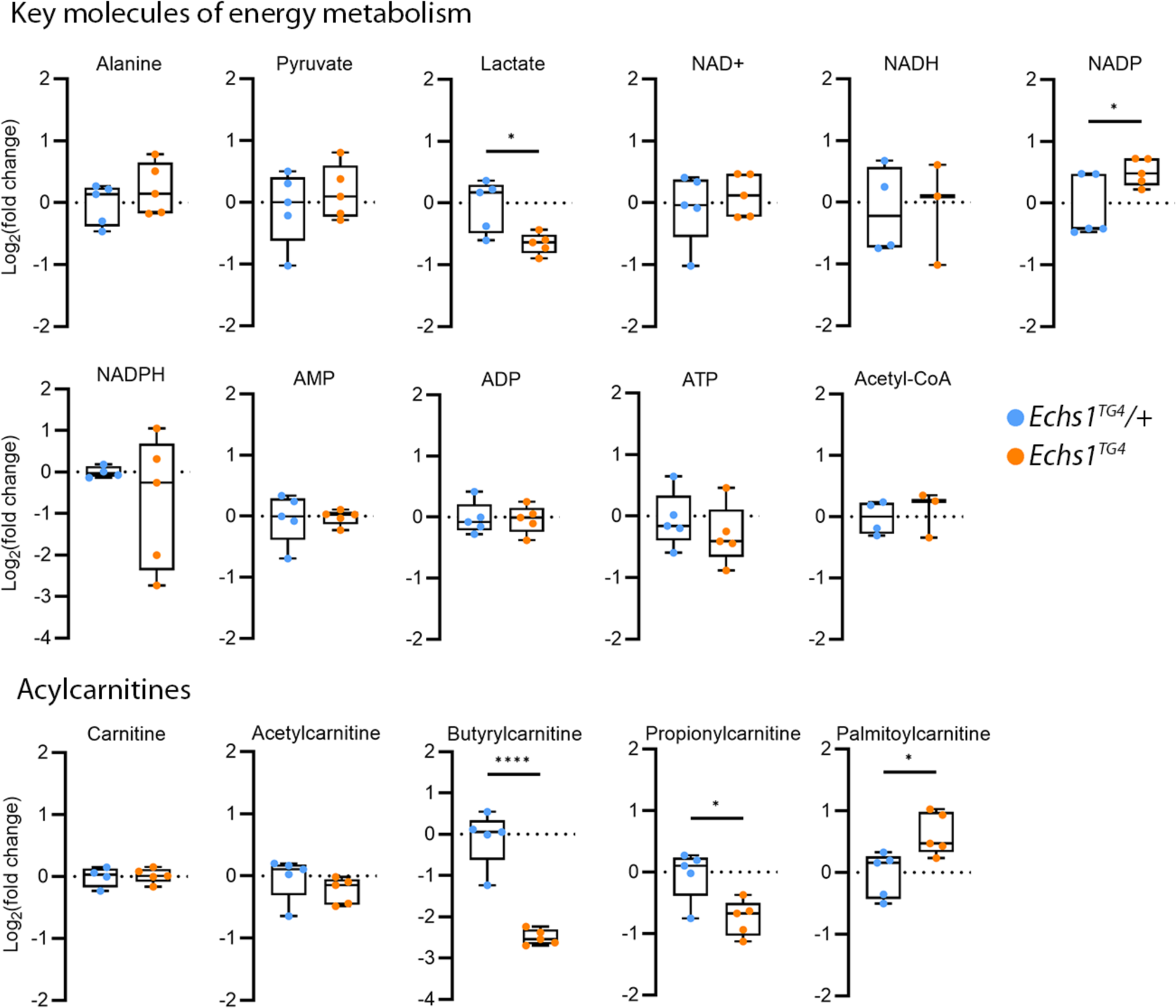
Central energy and fatty acid metabolism-related metabolite levels (Log2(fold change)) compared to control *Echs1^TG4^*/+ larvae.

### Tables

**Table S1.** Sugar yeast diet recipe.

| Sugar-Yeast diet |  |
| --- | --- |
| H<sub>2</sub>O | 1000 mL |
| Agar | 4.75 g |
| Dry yeast | 10.7 g |
| Glucose (dextrose) | 47.6 g |
| Raw sugar | 23.8 g |
| Semolina | 59.5 g |
| Propionic acid | 3.6 mL |
| Nipagin | 7.1 mL |
| Potassium tartrate | 7.1 g |
| Calcium chloride | 0.45 g |

**Table S2.**
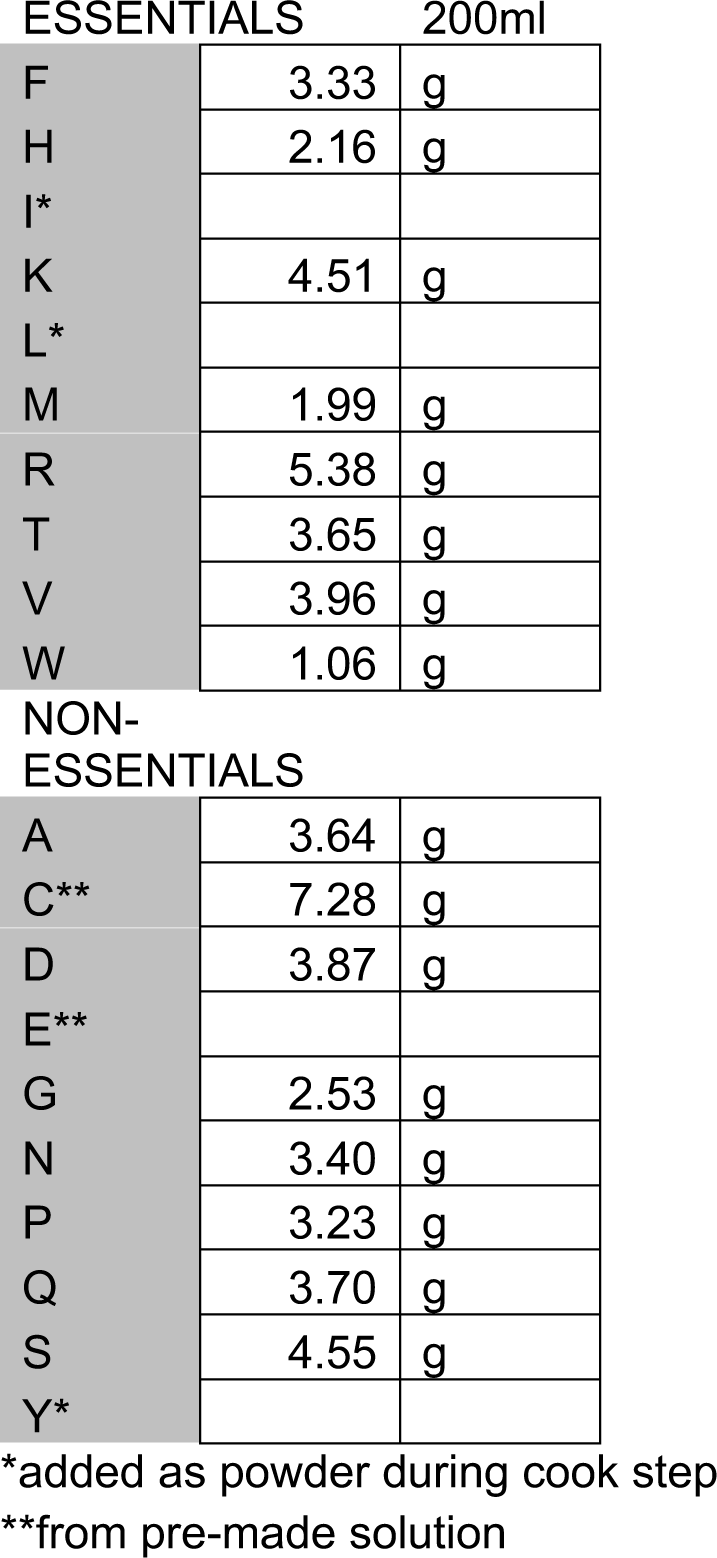
Holidic diet amino acid solution concentrations.

**Table S3.**
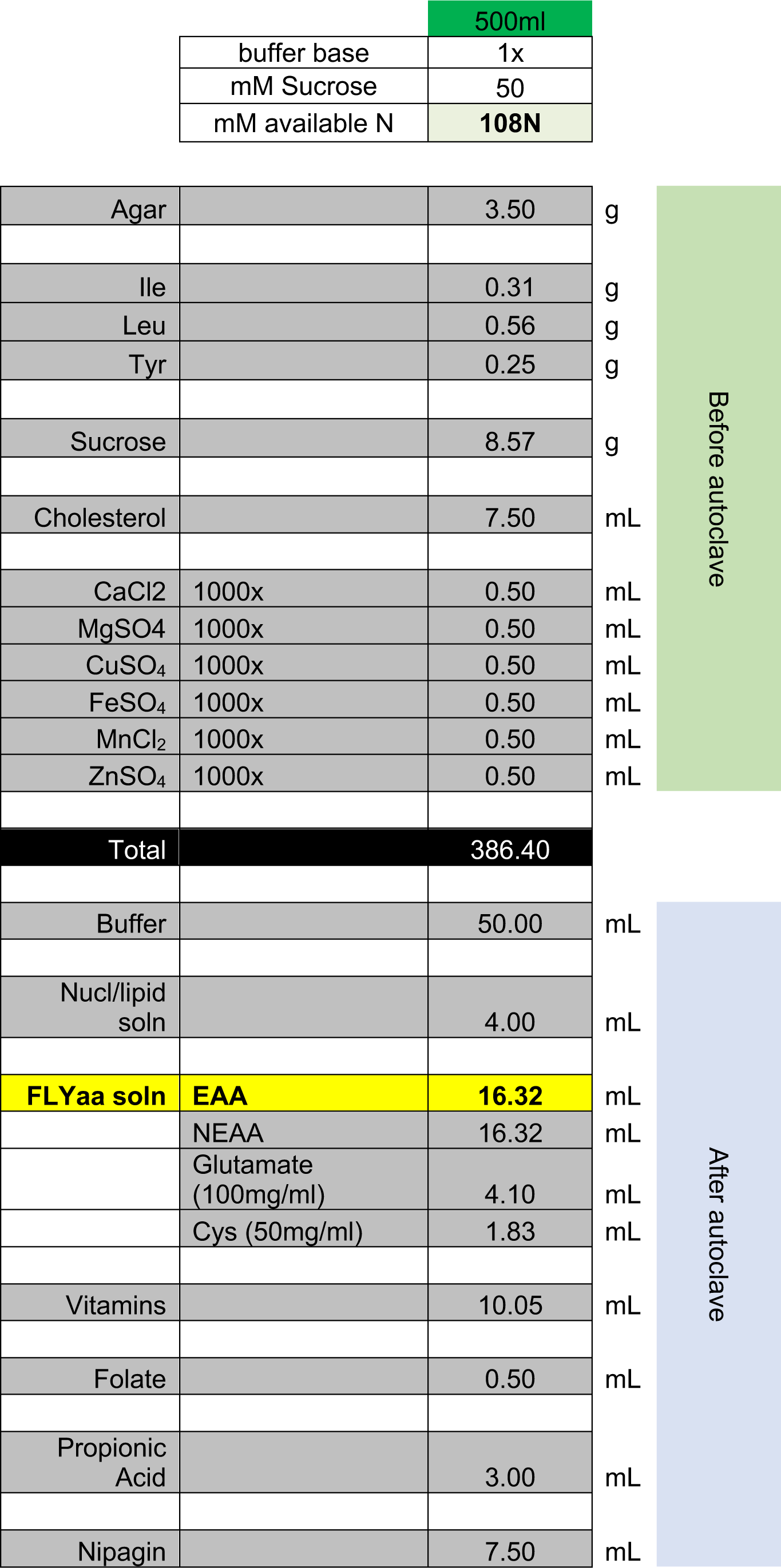
Holidic diet recipe.

**Table S4.** Specific Holidic diet modifications.

| <u>Valine restriction</u> |  |  |
| --- | --- | --- |
| Concentration | Valine in 100 mL * |  |
| 100% | 1.98 | g |
| 75% | 1.49 | g |
| 50% | 0.99 | g |
| 25% | 0.50 | g |
| *replaces valine amount in AA solutions table |  |  |

| <u>BCAA modification**</u> |  |  |  |
| --- | --- | --- | --- |
| Concentration | Leucine | Isoleucine | in 250 mL<br>Holidic cook |
| 150% | 0.42 | 0.23 | g |
| 125% | 0.35 | 0.19 | g |
| 100% | 0.28 | 0.15 | g |
| 75% | 0.21 | 0.11 | g |
| 50% | 0.14 | 0.08 | g |
| 25% | 0.07 | 0.04 | g |
| **valine added as part of AA solution |  |  |  |

